# Widespread false gene gains caused by duplication errors in genome assemblies

**DOI:** 10.1101/2021.04.09.438957

**Authors:** Byung June Ko, Chul Lee, Juwan Kim, Arang Rhie, DongAhn Yoo, Kerstin Howe, Jonathan Wood, Seoae Cho, Samara Brown, Giulio Formenti, Erich D. Jarvis, Heebal Kim

## Abstract

False duplications in genome assemblies lead to false biological conclusions. We quantified false duplications in previous genome assemblies and their new counterparts of the same species (platypus, zebra finch, Anna’s hummingbird) generated by the Vertebrate Genomes Project (VGP). Whole genome alignments revealed that 4 to 16% of the sequences were falsely duplicated in the previous assemblies, impacting hundreds to thousands of genes. These led to overestimated gene family expansions. The main source of the false duplications was heterotype duplications, where the haplotype sequences were more divergent than other parts of the genome leading the assembly algorithms to classify them as separate genes or genomic regions. A minor source was sequencing errors. Although present in a smaller proportion, we observed false duplications remaining in the VGP assemblies that can be identified and purged. This study highlights the need for more advanced assembly methods that better separates haplotypes and sequence errors, and the need for cautious analyses on gene gains.

## Introduction

Biological misinterpretations can occur when genomic regions unknowingly have errors. But it is unclear as to the magnitude of mis-assembly errors in existing genome assemblies, generated in the transition from the fragmented DNA sequences to the assembled blueprint of a species^1–8^. Followed by the first assembly of fruit fly in 2000^9^ and a human reference genome in 2003^10^, ∼100 reference genomes of vertebrates were deposited in public databases by 2010 using mostly intermediate read length (∼700 bp) Sanger reads. The number of genomes gradually increased to ∼700 by 2018, mostly using short read-based (∼35-250 bp) next generation sequencing (NGS)^11^. These genomes helped bring about discoveries in a variety of fields, including evolution, ecology, agriculture, and medicine^12–17^. However, with short read-based assemblies, it was difficult to resolve repeat regions longer than the read lengths^1,18–20^.

Preliminary studies have indicated that the longer the sequence read length, the less likely an assembly structural error^1^, which has been quantitatively validated in our companion Vertebrate Genomes Project (VGP) flagship study^5^. An underappreciated source of mis-assembly was heterozygosity^5^. Mis-assignment of heterozygous genomic regions led to both copies of the partnering alleles being assembled as paralogs in the same haploid assembly^3,5,6^, which are called false heterotype duplications by the VGP^5^. Likewise, accumulated sequence errors in reads, particularly long reads, led to under-collapsed sequences, which were called homotype false duplications^5^. Both heterotype and homotype false duplications in genic regions can be misinterpreted as gene gains^1,21,22^. The VGP proposed that these false gains happen in more highly divergent regions of the genome, where assembly algorithms have difficulty distinguishing haplotype homologs from haplotype paralogs^5^, but this was not quantitatively tested in regards to the type of duplication.

Although long-read sequencing is better at resolving repetitive regions^1,11,23^, they alone are unable to fully resolve false duplications^1,22,24^. One way to prevent false duplications is to make homozygous lineages through inbreeding. But this can be either impossible or very difficult under most circumstances^3,25^, especially if one were to sequence all species of a lineage, such as the goal of the VGP that aims to produce complete and error-free reference genomes for all ∼70,000 vertebrate species^26–28^. Another way to solve false duplications is to use assembly strategies for efficient haplotype phasing, some developed and applied in the VGP^5,24,29,30^. But most of the non-VGP vertebrate genomes in the public databases as of to date were reconstructed without haplotype phasing. A full quantitative and qualitative assessment has not been conducted on the prior versus VGP genomes to determine the extent and types of false duplications, and improvements in the VGP assemblies.

Here we performed a detailed analysis to measure the presence, magnitude and cause for false duplications in previous common reference assemblies and their VGP counterparts. We focused on three species, the platypus and zebra finch that were originally assembled using Sanger reads published in 2008^31^ and 2010^32^, and the Anna’s hummingbird that used short Illumina reads published in 2014^12,33^. These are popular references, with the associated studies collectively cited over 3,600 times as of April 2021 (Google Scholar). The VGP version of the assemblies were long-read based. We found widespread false duplications in previous assemblies that were corrected in the VGP assemblies, and also identified areas for improvement for future assemblies.

## Results

### Genome assemblies and identifying false duplications

The previous Sanger-based platypus^31^ and zebra finch^32^ reference genomes used standard pipelines for the best reference chromosomal level genomes at the time, generated with 500-1000 bp Sanger sequence reads, BAC-based scaffolding and FISH or cytogenetic chromosome mapping and assignments. No systematic effort was made for haplotype phasing but both the previous zebra finch and platypus assemblies were rigorously manually curated. The prior Illumina-based Anna’s hummingbird reference^12,33^ was generated with short reads (∼150 bp), and contigging and scaffolding with multiple paired-end and mate-pair libraries ranging from 200 bp to 20 kbp in size. An effort was made to remove alternate haplotypes during assembly. The VGP assemblies of the same species was generated with PacBio-based continuous long-read (CLR) contigs (N50 read length ∼17 kbp), which were scaffolded with 10X Genomics linked reads, Bionano Genomics optical maps, and Arima Genomics Hi-C chromatin interaction read pairs^5^. Systematic attempts to prevent false duplications were made, using FALCON Unzip to separate haplotypes after generation of contigs and purge haplotigs^34^ that search for and purge false heterotype duplications from the primary pseudo-haplotype assembly^5^. All VGP assemblies were subjected with rigorous manual curation to minimize assembly errors generated by algorithmic shortcomings. The previous and VGP assemblies of the zebra finch and Anna’s hummingbird were conducted on genomic DNA from the same individuals, and thus differences would only be due to sequencing platform and assembly methods.

The size of the previous assemblies of the zebra finch, hummingbird and platypus are 1.23 Gbp, 1.11 Gbp and 2.00 Gbp, respectively (**Supplementary Table 1**). They consisted of a total of 37,421, 54,736, and 958,970 scaffolds. Among the scaffolds, 35 and 19 super scaffolds were assigned to chromosomes for the zebra finch and platypus assemblies, respectively. The assemblies had 87,710, 70,084, and 243,835 gaps, and their average contig NG50 were 47.9, 27.0, and 11.4 kbp, respectively. The size of the VGP assemblies were all 0.05-0.17 Gbp smaller (**Supplementary Table 1**). They consisted of ∼280 to 3,140-fold fewer scaffolds (i.e. 135, 159, and 305 total), of which 33 (now 39 in our updated version), 33, and 31, respectively, were assigned to chromosomes, including the sex chromosomes. The number of gaps likewise were ∼160 to 470-fold lower, and contig NG50 were ∼250 to 1,320-fold higher: 12.0, 13.4 and 15.0 Mbp for the zebra finch, hummingbird and platypus, respectively. Alternate haplotype scaffolds of 0.95-1.58 Gbp in total size were separated from the primary assembly.

We performed self-alignment of each assembly using minimap2^35^ as a part of the purge_dups^30^ process to detect duplications independently from another assembly; purge_dups was created by members of the VGP in order to identify and purge false duplications in different contigs. Also, we aligned the previous assemblies to the new VGP assemblies of each species using the reference-free Cactus aligner^36^, which allows pair-wise detection of duplicates between the previous and new assemblies at the sequence and contig levels (**Fig. 1a,b**). We distinguished false duplications from true duplications, as we found that the former had read coverage at the haploid-level, gaps between duplications due to mis-assembly, and discordance in 10X linked-read pairs mapped back to the assembly. We classified each false duplication as heterotype duplications when heterozygous *k-mers* were found, and homotype duplications when read depth coverage was lower than the haploid-level, which occurs with sequence read errors, or when heterozygous *k-mers* were not found (**Fig. 1c**).

**Fig. 1.**
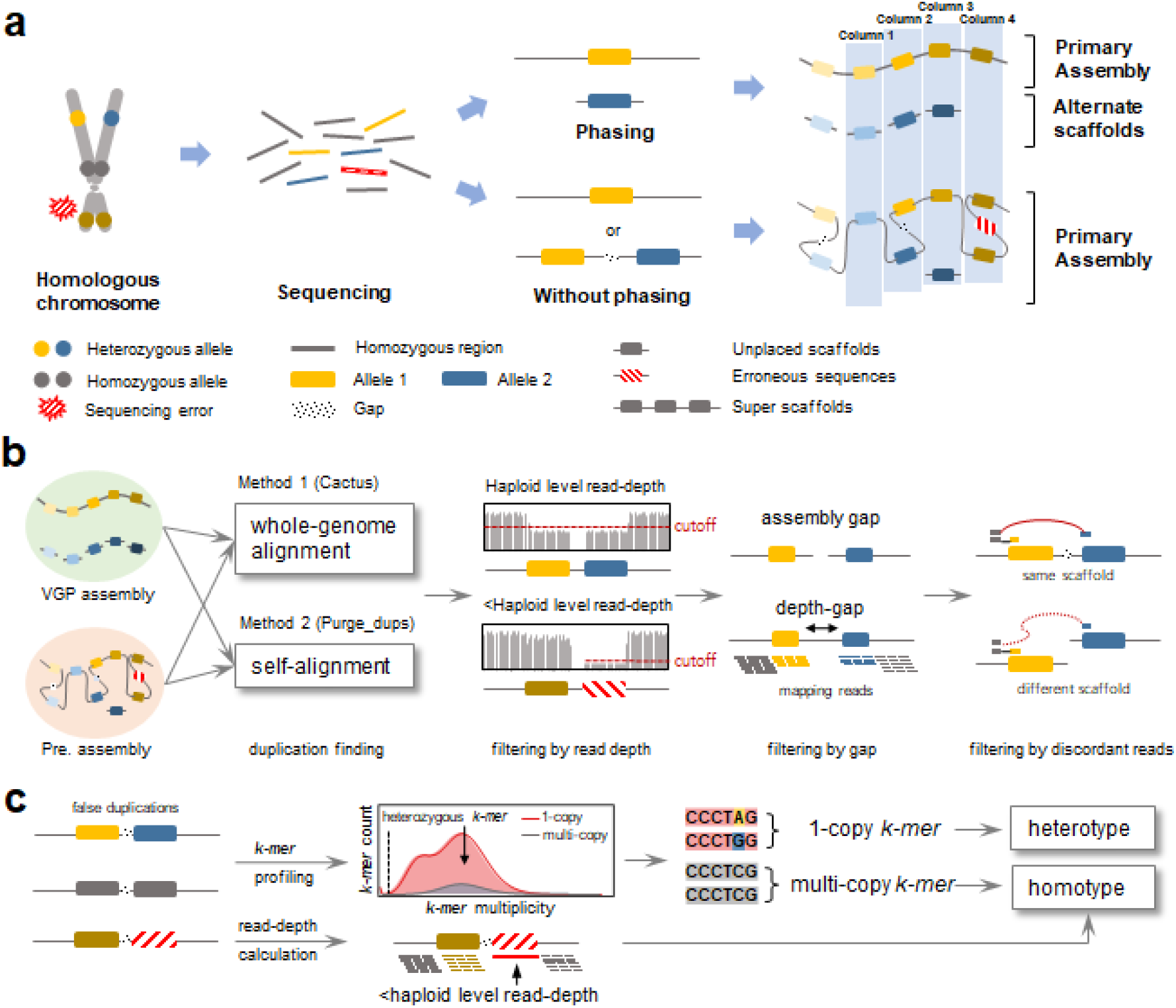
Overview to identify false duplication. **a**, Mechanisms of how false assembly duplications are created. If haplotype phasing is included and correctly done in the assembly process, there will be only one allele in the primary assembly, with the other either discarded or placed in the alternate assembly (Column 1). However, without proper phasing, both alleles of heterozygous loci may be assembled into one scaffold (Column 2) or two different scaffolds (Column 3) of the primary assembly. Alternatively, randomly or systematically piled up erroneous sequencing reads can generate false duplications (Column 4). **b**, Scheme to identify false duplications. Whole-genome alignment between the two assemblies using Cactus and self-alignment using purge_dups reveal candidate false duplicated regions or whole contigs. The union-set from these two independent methods is then used to find false duplications, which contain either below diploid read-depth of the 10X Genomics linked-reads, the presence of gaps between duplications, and discordance in read pairs between duplications. **c**, Scheme to classify false duplication types. Copy number and multiplicity of *k-mers* are calculated from the assembly and the 10X Genomics linked-reads respectively, and used to classify false duplications as heterotype or homotype. Heterotype duplication includes haploid specific *k-mers* (i.e. 1-copy). Homotype duplication does not include haploid specific *k-mers*, but does include sequencing errors that can be detected by read-depth below the haploid-level.

### False duplications in previous and VGP assemblies

The distributions of read depth coverage (**Extended Data Fig. 1**) and *k-mer* multiplicity (**Extended Data Fig. 2**) showed that previous assemblies included significant amounts of false duplications: 16% (196 Mbp), 4% (41 Mbp), and 6% (126 Mbp) of the total length of the prior zebra finch, Anna’s hummingbird, and platypus assemblies, respectively (**Fig. 2a, Table 1**). Heterotype was the major source of false duplication, an order of magnitude higher than the homotype except for the previous Anna’s hummingbird assembly (**Fig. 2a, Table 1**). Of the total false duplications, 7 to 24% were on the same scaffold, and the remainder on different scaffolds (**Table 1**).

**Fig. 2.**
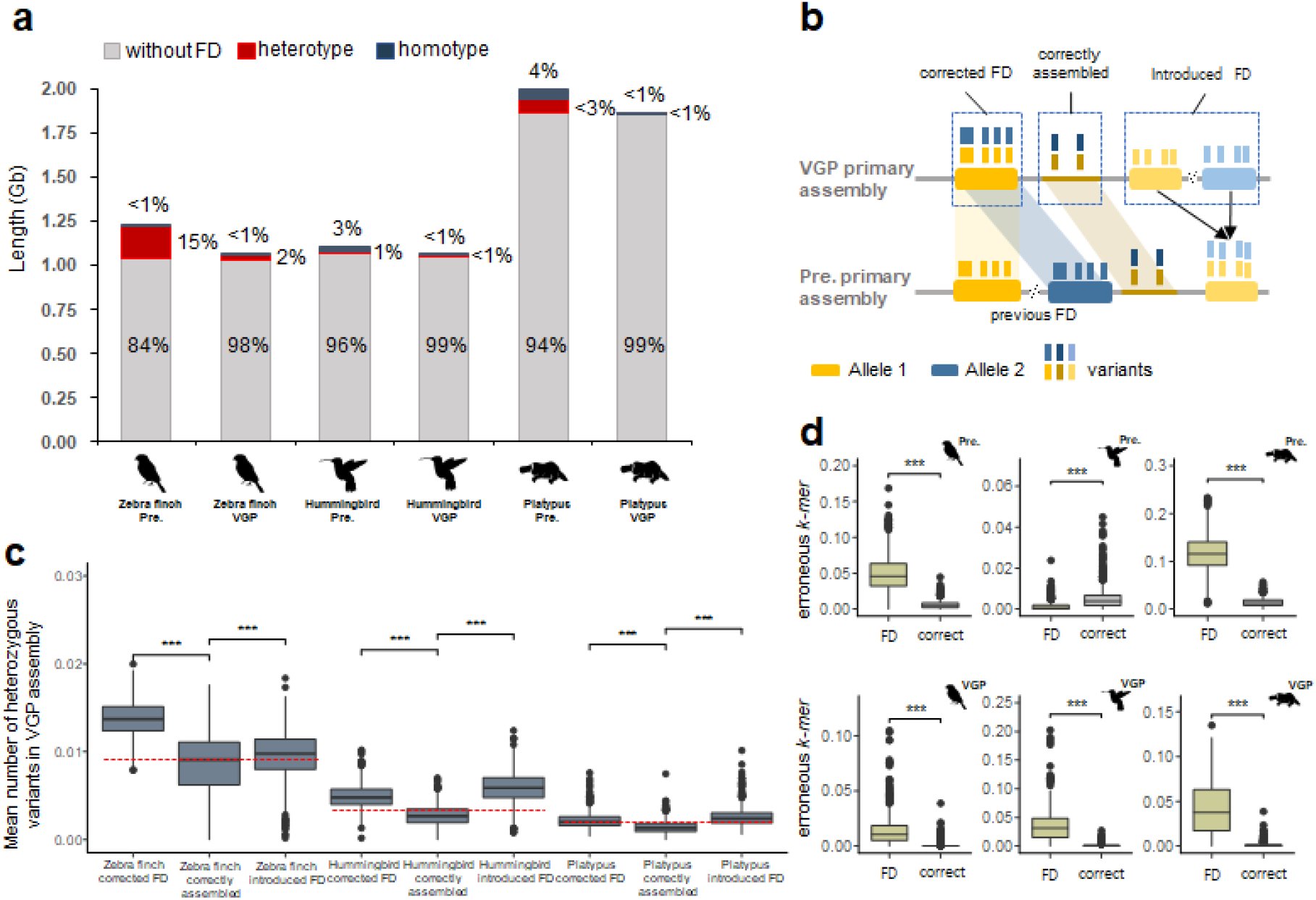
The amount of false duplication and factors that correlate with false duplication. **a**, The total assembly size and the proportion that are false duplications in the previous and VGP assemblies. False duplications were classified as heterotype and homotype. **b**, Scheme of false duplications (FD) in the previous and VGP assemblies due to heterozygous alleles. Corrected FD are regions in the VGP assembly that are false duplications in the previous assembly. Correctly assembled are regions without any false duplication in the previous and VGP assemblies. Introduced FD are false duplications introduced in the VGP assembly that were not present in the previous assembly. **c**, Heterozygosity of corrected FD, correctly assembled, and introduced FD, according to the VGP assembly haplotype data (\*\*\****P*** < 0.001; two-sided *T*-test). Red dotted line, overall heterozygosity of the genome. **d**, The portion of erroneous *k-mers* in false duplications and correct regions of each assembly (\*\*\****P*** < 0.001; two-sided *T*-test).

**Table 1.**
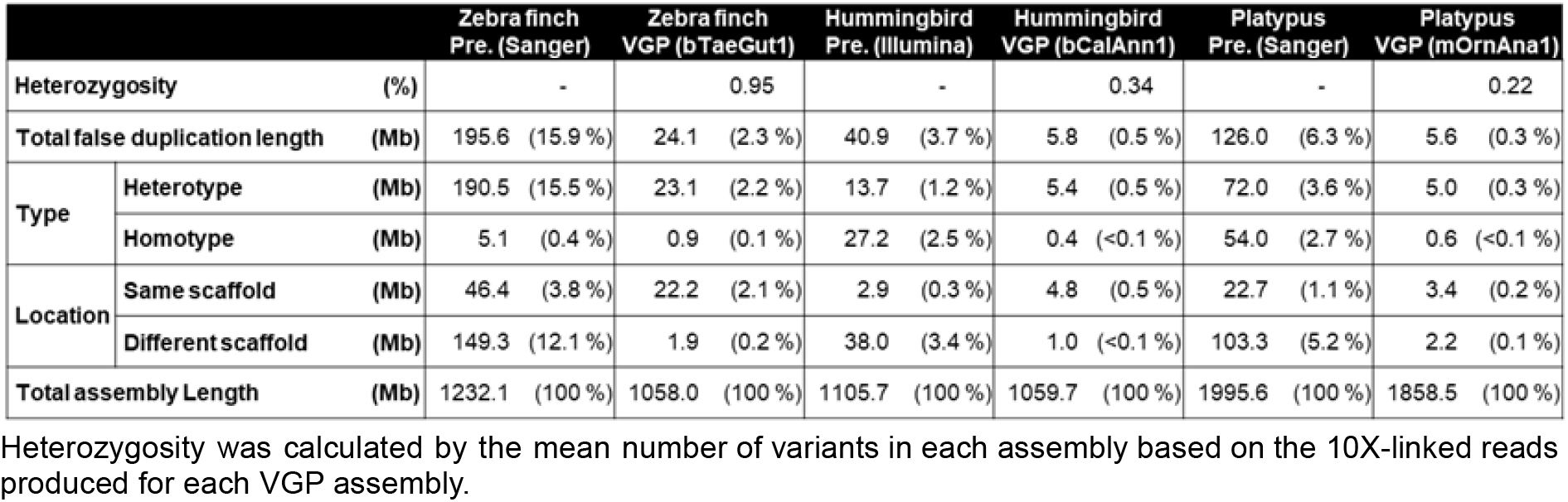
False duplication statistics in previous and VGP assemblies.

False duplications in the VGP assemblies were 7 to 22-fold less: 2.3% (24 Mbp), 0.5% (5.8 Mbp), and 0.3% (5.6 Mbp) of the total primary assembly in the zebra finch, hummingbird and platypus, respectively (**Fig. 2a, Table 1**). Heterotype was also the major type of false duplication. In contrast to the prior assemblies, there was a much higher proportion of the false duplications, 61-92%, found on the same scaffold, due to improved scaffolding using multiple long-range sequencing platforms.

### Heterozygosity and sequencing error on false duplication

The heterozygosity of false duplications in the previous assemblies that were corrected in the VGP assemblies (Corrected FD regions; **Fig. 2b**) were all ∼1.5 to 1.8-fold higher than correctly assembled regions without false duplications in both assemblies (*P* < 0.001; **Fig. 2c**). The heterozygosity of false duplications specific to the VGP assembly (Introduced FD regions; **Fig. 2b**) were all also higher (*P* < 0.001) with no specific level that differs with the previous assemblies (**Fig. 2c**). We also found more erroneous *k-mers* in false duplications than in the correctly assembled regions in both the previous and VGP assemblies (**Fig. 2d**). Further, regions between the false duplications were most of the time separated by an assembly gap and sometimes connected by unsupported sequence read depth gaps, due to incorrect gap filling or other assembly errors (**Fig. 3: Extended Data Fig. 3a,b,c**). These properties were not found for true duplications, including for a duplication of the acrosin (*ACR*) gene and an allele specific tandem duplication we found in the same contig with haploid level read depth of coverage (**Extended Data Fig. 4a,b**). These findings show that increased heterozygosity, especially those at the boundaries of homozygous and heterozygous sites, and sequencing errors are prone to false duplications.

**Fig. 3.**
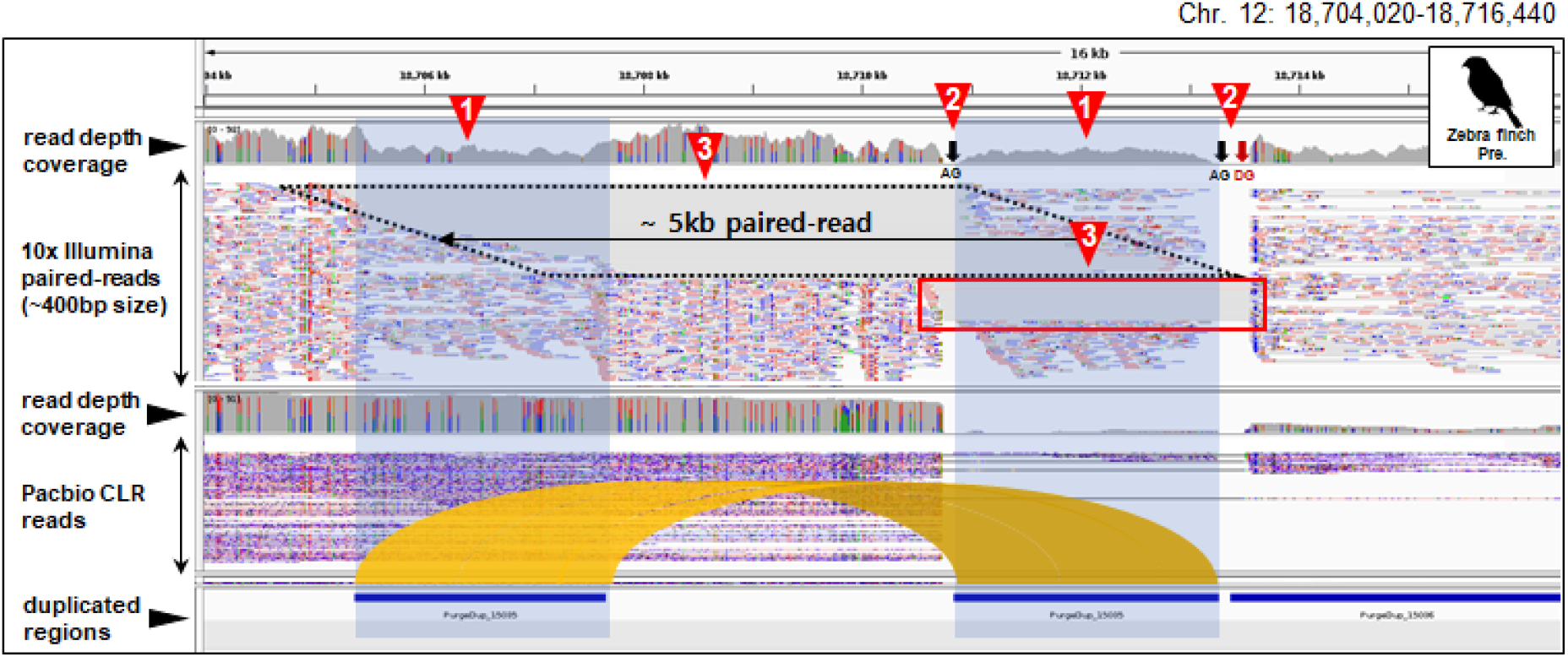
The presence of a gap and discordant reads between false duplications in the previous assembly. Shown is a locus in the previous zebra finch assembly with a false duplication. Characteristics are marked with triangles: 1) Nearly half depth-coverage and lack of heterozygous variants - colors indicating nucleotide heterozygosity; 2) Gaps between false duplications (black arrow); and 3) Discordant 10X linked reads (black dotted box). Red box, discordant reads found near the end of scaffolds that should be connected to each other. AG, assembly gap. DG, depth-gap (unsupported sequences by reads; see methods).

### False duplications cause false annotation errors

Among the false duplications, we found 4 to 24% of the coding genes were impacted in the previous assemblies, depending on species (**Fig. 4a**). Of these, we found three main types: 1) the majority being false gene gains [FGG] of nearly the entire coding sequence; 2) followed by false exon gains [FEG] within a gene; and 3) a minority being false chimeric gains [FCG] from a chimeric join among exons from different genes (**Fig. 4a,b; Supplementary Table 2**).

**Fig. 4.**
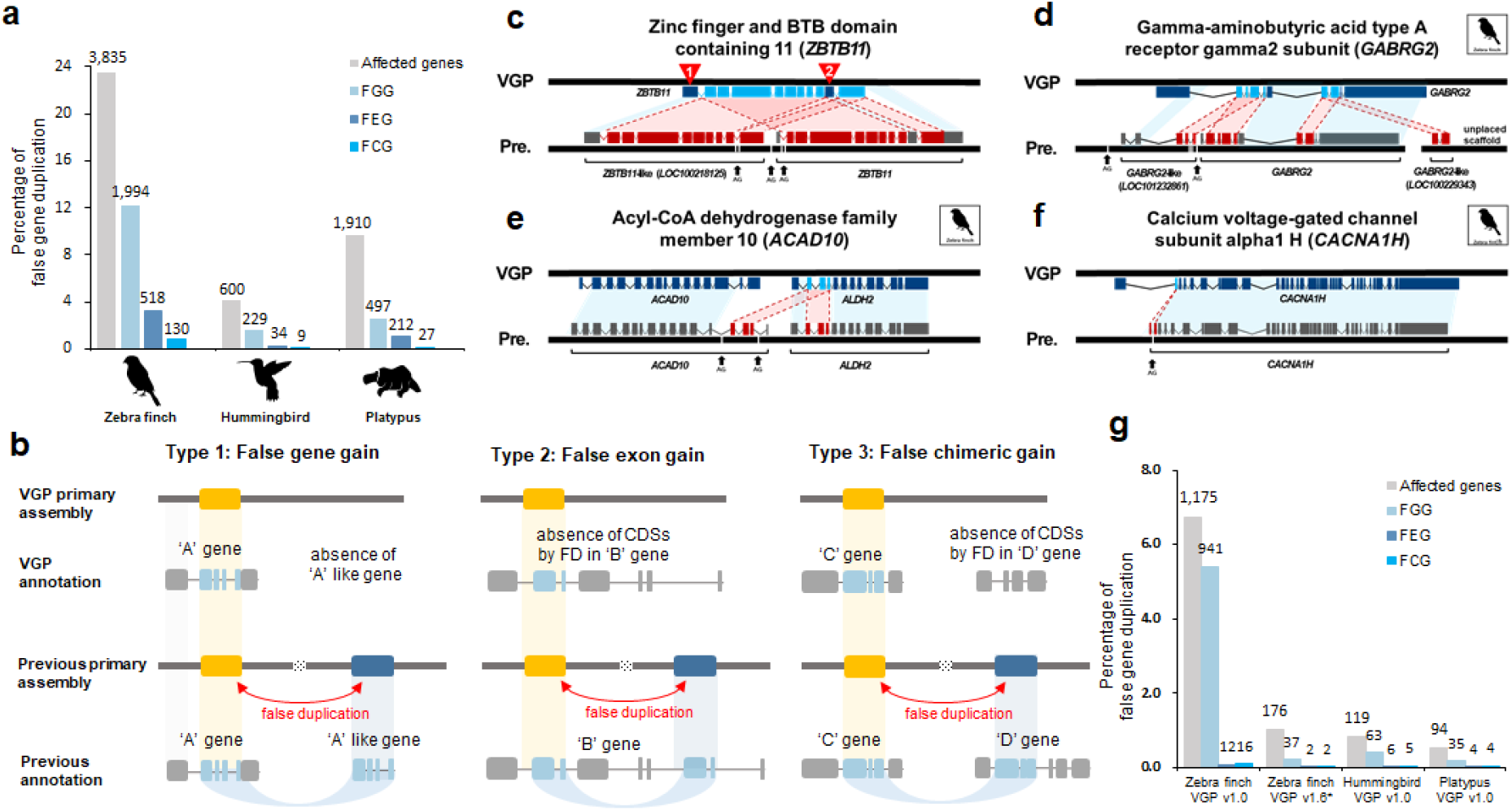
Mis-annotations due to false duplications. **a**, Amount and percentage of all genes with mis-annotations caused by false duplications in the previous assemblies. The amount of genes of each type is shown on the top of each bar graph. **b**, Types of mis-annotations caused by false duplications. When >50% of the CDS length of a gene was duplicated and annotated as another gene, and resulted in two genes with similar function (e.g. -like), we classified it to false gene gain (FGG, Type 1). When an exon within a gene was duplicated to itself, we classified it to false exon gain (FEG, Type 2). If the duplicated exon was inserted to another existing gene of different function, we classified it as a false chimeric gain (FCG, Type 3). **c**, FGG of *ZBTB11*. **d**, FGG of *GABRG2*. **e**, FCG involving *ACAD10* and *ALDH2*. **f**, FEG within *CACNA1H*. The red lines represent the connection between false duplications and the homologs in the VGP assembly. The blue boxes represent the homologous region between the VGP and previous assemblies. **g**, Amount of false gene annotation in VGP assemblies. Zebra finch VGP v1.0 was assembled with purging false duplications after scaffolding with purge haplotigs; zebra finch VGP v1.6 (* marked) was assembled with purging false duplications before scaffolding with purge_dups.

An example of a FGG included *ZBTB11* in the previous zebra finch assembly, was 9 of the 11 coding exons of *ZBTB11* falsely duplicated and annotated as *ZBTB11-like* (*LOC100218125*; **Fig. 4c**). The non-duplicated *ZBTB11* exon 1 (first exon) was included in *ZBTB11-like* and exon 2 (red mark; 10th exon) in *ZBTB11*, while these exons were assembled into one gene in the VGP assembly. The sequence alignment landscape of *ZBTB11* in the previous assembly showed typical characteristics of a false gene gain (**Fig. 5a**), whereas there was no sign of false duplications in the VGP assembly (**Fig. 5b**). The gamma-aminobutyric acid receptor subunit gamma 2 (*GABRG2*) was a complex example, where several false exon duplications were assembled in the same scaffold and annotated as a *GABRG2-like* (*LOC101232861*) FGG or as another *GABRG2-like (LOC100229343)* FGG on another scaffold, both with presumed false exon losses after the duplication from the original gene (**Fig. 4d**). Because true duplications can also be annotated as gene name-like, for example *ACR* and *ACR*-like (**Extended Data Fig. 4a**), the “like” term in the NCBI annotation can not be taken alone as evidence of a false duplication. *ALDH2* had three false duplicated exons that were incorporated into the adjacent *ACAD10* gene, causing a FCG for *ALDH2-ACAD10*, all with gaps around each of the false duplications (**Fig. 4e,5c**), none present in the VGP assembly (**Fig. 5d**). The calcium voltage-gated channel subunit alpha1 H (*CACNA1H*), a gene with specialized expression in vocal learning circuits of the zebra finch^37,38^, had a FEG in the second exon (**Fig. 4f**). Similar examples of FGG, FCG and FEG in the previous Anna’s hummingbird and platypus assemblies are shown in **Extended Data Fig. 5**. This includes false duplications that overlap in the CDSs of *ATF3, PCBD1* and *VAMP4* in the previous hummingbird assembly; and of *ZP2, UPF2* and *HSF2* in the previous platypus assembly. The platypus vomeronasal receptors (*V1R*) gene family expansion was reported as a sensory adaptation for underwater life history^31^; our results show 43 of the 267 annotated *V1R* genes (16%) are actually false duplications in the previous assembly (**Supplementary Table 3**).

**Fig. 5.**
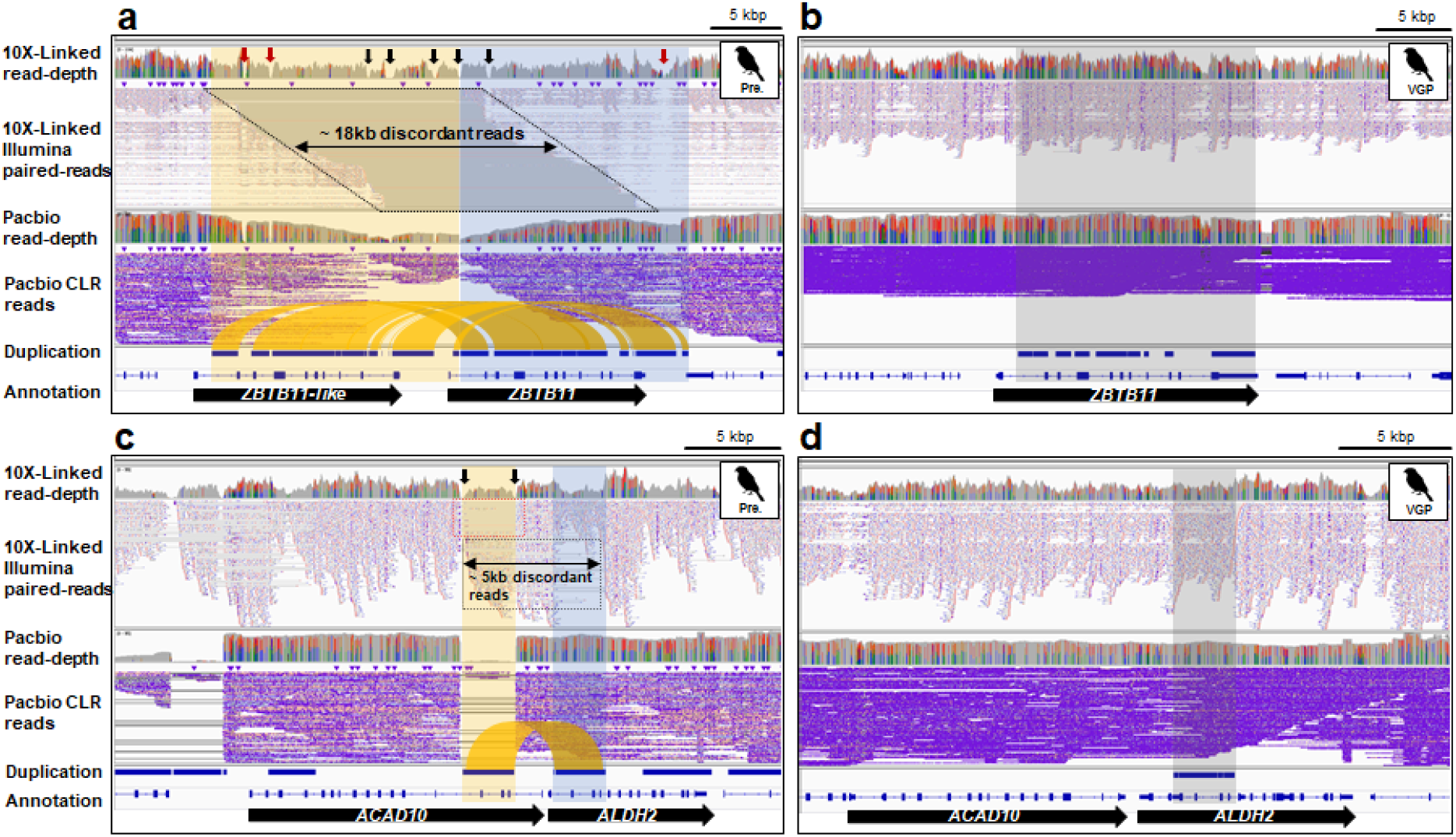
The genome landscape of false gene gains. **a**, The genome landscape of *ZBTB11* and false *ZBTB11*-like (*LOC100218125*) genes in the previous zebra finch assembly. Most of the region of *ZBTB11* was duplicated adjacent to itself in the previous assembly (highlighted as orange and blue) and show typical characteristics of false heterotype duplications. **b**, Corrected gene structure in the VGP assembly (grey). **c**, The genome landscape of *ACAD10* and *ALDH2* genes in the previous zebra finch assembly. Three exons of *ALDH2* were inserted in *ACAD10* by a false duplication (highlighted as orange and blue). **d**, Corrected gene structure in the VGP assembly. The extrinsic three exons from *ALDH2* (grey) were not found in *ACAD10* of the VGP assembly. The different colors and their heights in the read depth rows are the proportion of sites in reads with haplotype variants.

Among non-coding sequences, long terminal repeats (LTRs) sequences of the zebra finch were reported to have expanded 2.5 times more than chicken^32,39^ and short interspersed nuclear elements (SINEs) were reported to be highly expanded in the platypus relative to other mammals^31^. However, we found 18,757 copies of LTRs (21% of the total) were with false duplications in the previous zebra finch assembly and 140,279 copies of SINEs (6.1% of the total) were false duplications in the previous platypus assembly (**Supplementary Table 4**). In the previous Anna’s hummingbird assembly, 3 to 5% of LTRs, SINEs, and long interspersed nuclear elements (LINEs) were false duplications (**Supplementary Table 4**).

### False duplication and annotation errors remaining in VGP assemblies

The VGP assemblies used in this study were produced with the VGP v1.0 pipeline, where heterotype duplications were removed by purge haplotigs after scaffolding. In addition, many were detected and removed during manual curation^5^. Although the amount of false duplication in VGP assemblies was drastically lower than previous assemblies, here we found 74 to 119 scaffolds included some false duplications, of which 5 to 34 (3-11% of the total number of scaffolds) were completely duplicated (**Fig. 6**). From this error, we observed 1,175, 119, and 94 genes of the zebra finch, hummingbird and platypus were total or partial false duplications in the VGP v1.0 pipeline (**Fig. 4g**). False duplications were observed within both named chromosomes and unplaced scaffolds, with no discernable patterns in terms of chromosome (**Extended Data Fig. 6a,c,e**). However, for some small unplaced scaffolds (< 50 kbp) the proportion of their scaffolds as false duplications were large, with some cases where the entire scaffold was a false duplication (**Extended Data Fig. 6b,d,f**). This indicates that for the VGP assemblies, some unplaced scaffolds are simply the other haplotype or homotype duplication, and are within the range of long-read lengths (1 to 50 kbp).

**Fig. 6.**
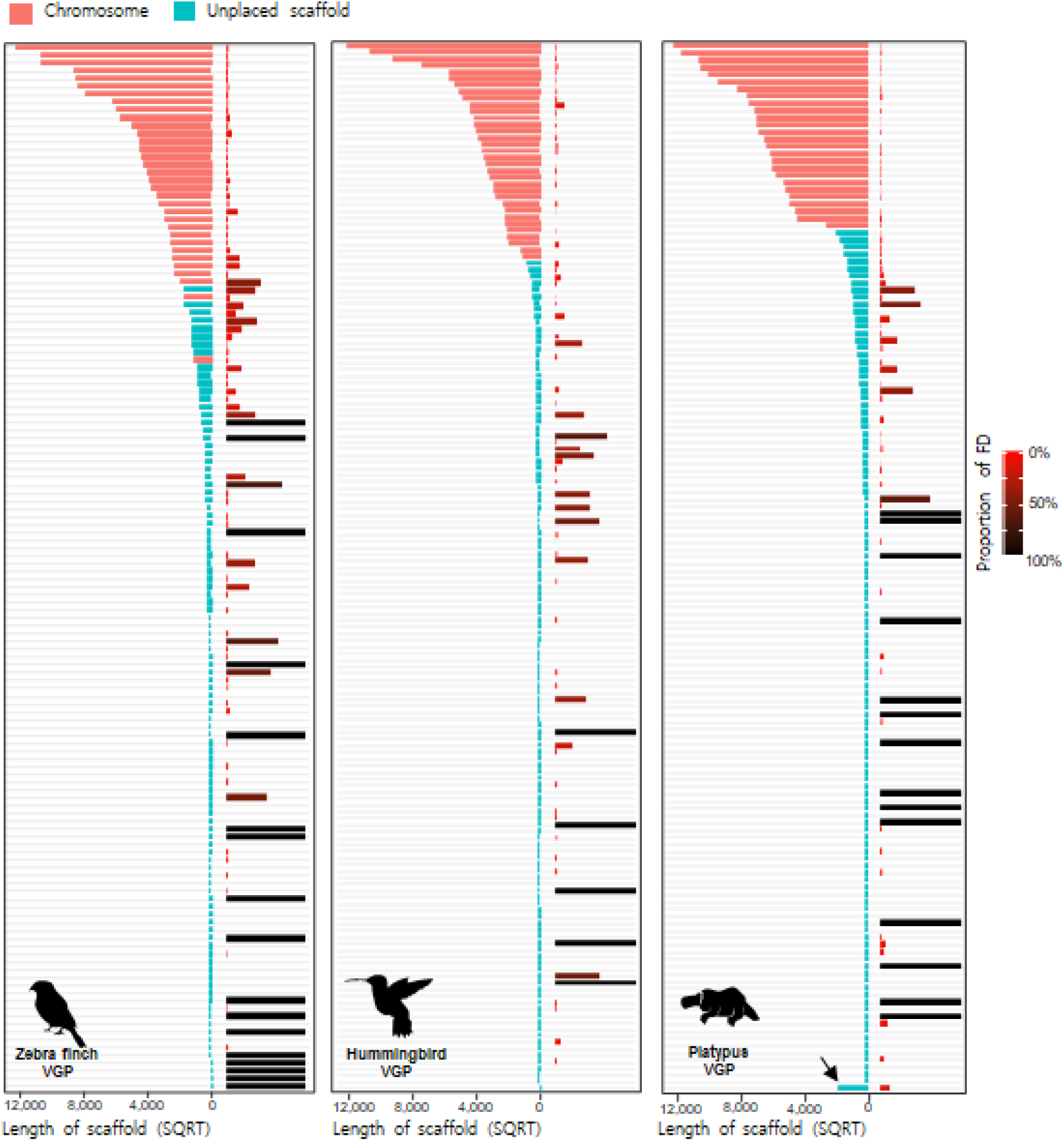
False duplications left in VGP assemblies. The left side of each graph shows the scaffold length of named chromosomes (pink) and unplaced scaffolds (turquoise). The right side shows the proportion of each scaffold that is falsely duplicated either within the same or different scaffolds. Arrow: for the platypus, scaffolds < 40 kbp scaffolds were concatenated into the one scaffold, where we found 20 scaffolds were completely duplicated among them.

We manually verified examples, and found some of the same type of errors seen in the previous assemblies, except the duplications were larger, presumably due to the longer read lengths and long optical maps of the VGP assemblies. An example was a false gene gain of *NPNT*, called *NPNT-like* (*LOC100218132*), on chromosome 4, named as such by the NCBI annotation pipeline applied to the VGP zebra finch 1.0 assembly (**Fig. 7a**). However, the false duplication structure caused 4 missing exons in the 5’ region of *NPNT* and 3 missing exons in the 3’ region of *NPNT-like*. Characteristic of the prior assembly, the false duplications were separated by an assembly gap, with discordant 10X linked-reads and at haploid depth coverage. Other examples included those that contained non-coding sequence (**Extended Data Fig. 7a**), and those were false chimeric Pacbio palindromic reads that lead to false duplications of the read length (7-17 kbp), with 10X linked-read depth gaps (**Extended Data Fig. 7b,c**). A case of large duplications was on zebra finch chromosome 29, where 4 segments adding up to ∼1.9 Mbp total were classified as false duplications using our criterion, making up ∼45% of the assembled 4.2 Mbp microchromosome (**Fig. 7b**).

**Fig. 7.**
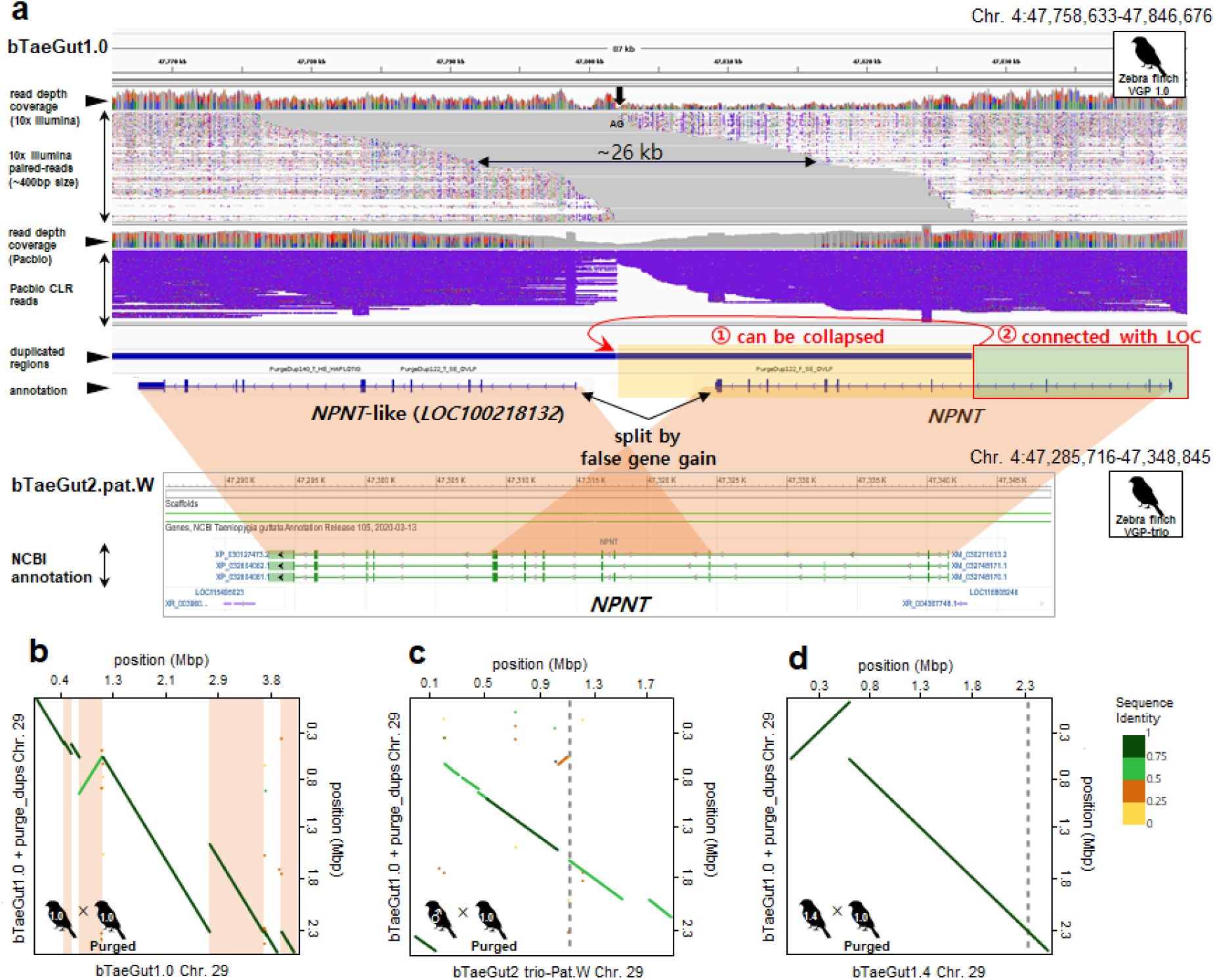
Haplotype variation results in a false gene split and gene gain in the VGP assembly. **a**, The *NPNT* gene in the VGP zebra finch assembly bTaeGut1.0 (first release) has *NPNT*-like gene adjacent to it with an assembly gap (AG) and discordant 10X linked-reads in this region. In contrast, the trio-based assembly (bTaeGut2.pat.W) had no *NPNT*-like gene, suggesting a false gene gain in bTaeGut1.0. The false duplication we found in this region was collapsed by purge_dups, and the falsely segmented gene structure was recovered. The VGP assembly v1.7 pipeline with purge_dups conducted before scaffolding prevented this false duplication (**Extended data Fig. 8**). **b**, Dot plot of alignment showing large ∼1.9 Mb false duplication of chromosome 29 (apricot) in the zebra finch VGP v1.0 pipeline assembly, bTaeGut1.0. **c**, The large ∼1.9 Mb of duplications of chromosome 29 in bTaeGut1.0 were prevented in the trio-based assembly. **d**, The 1.8 Mb duplication was prevented with purging pre-scaffolding in the VGP v1.7 pipeline. After purging pre-scaffolding, the false duplications in this study seen in bTaeGut1.0 were prevented in bTaeGut1.4. The boundaries of the scaffolds are represented as grey dashed lines.

To verify whether many of these false duplications are due to false haplotype separation, we examined a VGP trio based assembly of another zebra finch individual^5^. This trio based approach was recently developed with the goal of using parental short reads to separate out haplotype sequences of the child long reads, before contig assembly and scaffolding^5,24^. We found that both the local *NPNT* (**Fig. 7a**) and the large ∼1.8 Mb of duplications of chromosome 29 were prevented in the trio-based assembly (**Fig. 7c**).

We sought a means to further prevent false duplications in a non-trio based VGP assembly, as the individuals used in this study do not have available parental data. As done for some later assemblies of other species^5^, but not directly tested on the same individual, we reassembled the zebra finch individual originally used as the reference, but performed purge_dups before scaffolding contigs in the VGP v1.6 pipeline rather than afterwards in the VGP v1.0 pipeline. We also added a new tool called Merfin, to polish the assembly with long reads (https://github.com/arangrhie/merfin), a step that does not influence false duplications but improves base level accuracy. We called the update VGP v1.7 pipeline. After reassembly, the *NPNT* and other false duplications were prevented (**Extended Data Fig. 8, Supplementary Table 5**). The 1.8 Mb of false duplications on chromosome 29 were prevented, resulting in chromosome size and alignment consistent with the size of chromosome 29 after identifying false duplications found in this study on the original VGP 1.0 pipeline assembly (**Fig. 7d**). Overall, we observed a reduction from 1,175 genes to 176 genes with false duplications (**Fig. 4g**), and reduction of 16 entire duplicated scaffolds to 5 (**Supplementary Table 5**). These findings show that false duplications are still prevalent in some of the best assemblies, but have potential to be removed with improved haplotype phasing.

## Discussion

In this study, VGP assemblies^5^ were compared with previous assemblies of the same species/specimen to quantify the potential of false duplications. We found duplicated regions in the previous assemblies that showed characteristics of false duplications, which we quantified across the entire assemblies here for the first time. Characteristics we find that can be used to determine whether a gene of interest or entire genic regions is a false duplication include: 1) half depth coverage for each duplication for heterotype duplications, or very low depth coverage on one duplication for homotype duplications; 2) presence of gaps between duplicated pairs on the same scaffold; 3) discordant or spanned linked read pairs used for scaffolding, whenever 10X or other types of paired reads of a DNA fragment; and 4) 1 copy *k-mers* for heterotype duplications. Some of these characteristics have been reported in other studies prior to the VGP effort^5,21^, but not in a systematic manner of comparing previous and new assemblies that attempted to remove false duplications as reported here. The false duplications were highest in the previous Sanger-based assemblies and lowest in the VGP Pacbio-based long read assemblies that purged them before scaffolding or in a VGP trio Pacbio-based long read assembly that sorted haplotype reads before contig and scaffold generation. One major source of the false duplications was a near doubling in the level of heterozygosity in the false duplicated regions compared to the rest of the genome. Further, the species with the highest heterozygosity, the zebra finch, had the highest proportion of false duplications in the previous and VGP assemblies. Another major source, sequencing error, was also found in false duplicated regions in both previous and VGP assemblies.

These false duplications led to mis-annotations as false gene, exon, and chimeric gene gains. When the duplication is created, the inserted allelic sequence results in annotation of two similar genes. These types of false gain errors were made in genes involved in important phenotypes, leading to serious misinterpretations in downstream analysis^40^. For example, false gene gains reduce one-to-one orthologs, which are key genes used in comparative genomics and phylogeny. When false gain errors occur in an expanded gene family of closely related genes, this leads to false positive cases of gene family expansions as we report here, others previously^41^, and in a companion study on the oxytocin family of receptors^42^. For phylogeny, these duplications create false orthologs or indels in genes that weaken gene- and species-inferred relationships. This can be made worse with multiple false duplications of genes with closely related paralogs, such as the overestimated LTR expansion in the zebra finch^32,39^ that we find here. Our findings indicate that caution should be taken when interpreting gene family expansion in assemblies generated without haplotype phasing and checking for false duplications, using the approaches we describe here.

Our findings that heterotype false duplications are much higher than homotypes, indicates that proper haplotype separation is still a current problem in genome assembly, even when they have been greatly reduced in the VGP assemblies. The VGP 1.6 trio pipeline removes more heterotype false duplications^5^, but it requires parental sequence data to sort haplotypes, and parents will not be available for all individuals. Scanning regions around gaps with reads and *k-mer* profiling, and discordantly mapped linked reads or disconnected Pacbio reads should be helpful in identifying false duplications in any assembly. However, the best way to prevent these we propose would be to improve haplotype phasing of raw reads without parental data, remove reads with sequence errors before assembly, and generate complete diploid genome assemblies.

The VGP group is constantly updating its sequencing and assembly pipeline to create a genuine blueprint for assembly of complex and large genomes as found among vertebrates. Doing so requires in depth evaluation of assemblies, as done in this study. In the VGP assembly pipeline, the continuous long read (CLR) data type of Pacbio sequencing is being replaced by the closed circular sequence (CCS) high fidelity (HiFi) read data type^5,43^, which reduces the base-pair error rate without the need for short-read Illumina polishing. We expect these new HiFi reads to also reduce the false duplications due to sequence errors, and it may allow better separation of haplotypes. The sequence read lengths, however, are currently ∼20% shorter (15-20 kbp), and thus may lead to less contiguity across real duplications longer than the read lengths. Our findings emphasize that creating error-free reference genome assemblies should be a fundamental requirement of future genomics and biology.

## Acknowledgement

We thank the members of the VGP assembly group for feedback and support. We thank David Clayton, Christopher Balakrishnan, Julia George, Mahalia Frank, and Katy Palios for pointing out to us the large duplications in zebra finch chromosome 29 and several other chromosomes of the VGP 1.0 assembly. We thank Dengfeng Guan for modifying purge_dups for this research.This study was supported by the Marine Biotechnology Program of the Korea Institute of Marine Science and Technology Promotion (KIMST) funded by the Ministry of Ocean and Fisheries (MOF) (No. 20180430), Republic of Korea to HK, and the Howard Hughes Medical Institute (HHMI) to EDJ.

## Materials and Methods

### Assemblies and read data

The primary assembly of the previous and VGP version of the male zebra finch, female Anna’s hummingbird, and female and male platypus were downloaded from NCBI by ftp along with their assembly statistics, gaps, repeats and annotation data (**Supplementary Table 1**). For the VGP assemblies, we included both the primary and alternate pseudo-haplotype sequences. The raw reads used for the previous assemblies of the zebra finch and platypus generated by Sanger sequencing were not available to download from the sequencing read archives (SRA) on NCBI. The PacBio CLR and 10X raw reads used to generate the VGP assemblies were downloaded from the VGP Genome Ark (https://vgp.github.io/genomeark/).

### Code availability

All source codes used for identifying false duplications and false gene gains are available in github platform https://github.com/KoByungJune/FalseDuplication.

### Identifying false duplications

#### Candidate duplications from sequence similarity

We identified false duplication candidates by sequence similarity in whole genome alignments between the previous and VGP assemblies and self-alignment of an assembly to itself. We used Cactus^36,44^ to generate whole genome alignment across assemblies and HAL^45^ to transform the Cactus results into a readable multiple alignment format. One to many homologs between two assemblies of the same species were then considered as potential false duplication candidates. Since the Cactus alignment contained very short sequences (<20 bp) in alignment blocks, we filtered out blocks shorter than 20 bp or query sequence coverage of less than 80% to avoid spurious alignment noise. Self-alignment was performed with minimap2^35^ ‘-xasm5’ option on, after segmenting contigs by ‘N’-base gaps. Purge_dups was then used to find false duplications^30^. We used a purge_dups version that we asked the developers to modify (‘add_loc’ branch in github of purge_dups) to output the pair-wise homologous loci for each false duplication found.

#### Filtering true duplications

False duplication candidates were further distinguished from true haplotype specific duplications using 10X linked-read alignments; it was difficult to map PacBio CLR reads to the previous assemblies, as the length of the majority of the contigs of the prior assemblies (e.g. 1∼3 kbp) were shorter than a read length of the VGP assemblies (e.g. ∼10-17 kbp). The paired-end reads from the linked reads were aligned with EMA v0.6.2^46^ to keep the barcodes, and BWA v0.7.17^47^ without the barcodes. Coverage distribution across the entire assembly was extracted using samtools^48^. False duplication candidates from purge_dups self-alignment were further processed using the rest of the purge_dups pipeline, to finalize false duplications using this coverage distribution. Candidates from the Cactus alignments were similarly filtered using the same read depth threshold as in purge_dups. Any duplications with lower than half the diploid read depth of coverage were further considered. We then applied two additional criteria: 1) presence of scaffolding gap or read depth-gap between a duplication pair; and 2) discordant read pair alignments. A depth-gap is defined as a region with no read alignments between duplication pairs, which occurs from incorrect gap-filling or incorporation of reads with sequencing errors (**Extended Data Fig. 3**) during assembly. A discordant read pair was defined when the insert size between the pairs is unexpectedly large (>550 bp) or mapped to another scaffold as in Kelley and Salzberg^3^. We required both presence of discordant reads and concordant reads to align, where one end from a discordant read pair and concordant read pair aligns to the identical flanking region (∼550 bp) of a duplication, while the other end aligns to each of the homologous duplications.

#### Classifying heterotype and homotype duplications

The filtered false duplications were further classified based on *k-mer* (*k* = 20) analyses. We extracted 20-mers from the assemblies and 10X linked-reads using Meryl^22^ and performed Merqury^22^ analysis to obtain the *k-mer* spectrums. Using the *k-mer* spectrum, we defined erroneous *k-mers* as those found in the assembly with read multiplicity lower than 6, 3, and 18 in the previous assemblies of zebra finch, hummingbird, and the platypus, and 3, 3 and 10 for VGP assemblies, respectively. Likewise, any non-erroneous *k-mer* found once in the assembly was defined as a single-copy *k-mer*. We classified false duplications as heterotype duplication when both of the duplicated pairs had single-copy *k-mers* with average read depth over than sequencing error, 5, 8 and 22 for the previous assemblies of zebra finch, hummingbird, and the platypus, and 2, 2, 9 for the VGP assemblies; otherwise as homotype duplication, which had no single-copy *k-mer* found on either side of the duplication or one duplication of the pair had read depth below heterotype duplication levels.

### Evaluating false duplications

PacBio CLR reads were mapped to both the previous and VGP assemblies using minimap2^35^. The mapped reads on each assembly were visualized with IGV^**49**^. Duplications found in the VGP assemblies were aligned to its counterpart assembly and visualized with D-Genies^50^. The location of false duplications in VGP assemblies was visualized by karyoploteR^51^.

The heterozygosity of assemblies and each corrected FD and correctly assembled region was calculated as the number of variants divided by the length of the region, with 1,000 bootstrapping replicates to generate a distribution used in a Student’s *t*-test between those regions. To calculate heterozygosity in the region of the introduced FD in the VGP assemblies, we masked false duplications as ‘N’s, then the variant was estimated from newly mapped 10X linked-reads onto the masked assembly, followed by the same bootstrapping and statistical approach as used above. Samtools and bcftools were used for variant calling with the multiallelic model. The sequence error rate of each duplicated and correct region was calculated by dividing the number of erroneous *k-mers* by the total number of *k-mers* found. The distributions of sequencing error rate for duplicated and correct regions were also generated by 1,000 bootstrapping replicates, and a Student’s *t*-test was performed on those distributions.

### Identification of false gene gain annotation errors

We calculated the number of genes affected by false duplication, with the later defined as regions with duplicated sequences that overlapped with the CDS regions of an assembly. The Refseq annotation of NCBI was used and only the longest CDS of all isoforms generated from each gene was used. The genes influenced by false duplications were classified into three types: 1) false gene gain (FGG) in which a gene was falsely duplicated almost entirely or partially over 50% of the CDS length; 2) false exon gain (FEG) of one or more exons within the same gene; and 3) false chimeric gain (FCG) in which duplicated exons from one gene were inserted into another gene. FGG, FEG, and FCG were included only when at least one coding exon of a gene completely overlapped the false duplication. To visualize the example cases of mis-annotation, GSDS 2.0^52^ was used.

To search for possible false duplications overlapping with non-coding repetitive elements, we counted the number of LTRs, SINEs, and LINEs affected by false duplications using NCBI repeat information generated by repeatMasker^53^.

In the platypus, we also searched false segmental duplication and false gene gains of the *V1R* family in the same manner as above. We used segmental duplication DB from http://eichlerlab.gs.washington.edu/database.html ^31^. We checked for 267 *V1R* genes for potential false gene gains in the previous assembly of the platypus, which included “ORNANAV1R” in the symbol of the gene. We also identified *V1R* genes in VGP assembly using the same method.

### False duplication correction in zebra finch assembly using the VGP pipeline v1.7

We reassembled the zebra finch assembly using a variation of the VGP v1.6 pipeline, we called VGP pipeline v1.7. Aside from software updates, the two main differences with respect to VGP pipeline v1.0^5^ are: 1) purge haplotigs was replaced by purge_dups, for more effective purging of false haplotype and homotype duplication; 2) during the final Arrow polishing step variant calls were filtered with Merfin (https://github.com/arangrhie/merfin), to avoid introducing erroneous *k*-mers in the assembly. This resulted in the following assembly steps: 1) FALCON unzip contig assembly; 2) purge_dups to purge false duplications in the primary assembly, and place them in the alternate assembly; 3) scaffolding the primary assembly with 10X linked reads and scaff10X software; 4) scaffolding with Bionano optical maps and Bionano solve software; 5) scaffolding with Arima Genomics Hi-C and Salsa v2.2 software; 6) polishing with the long reads using Arrow and filtering the variant calls with Merfin; and 7) a final polishing with longranger aligner and freebayes. We added the assembled mitochondrial genome assembly prior to the polishing steps to prevent overpolishing of NUMTS in the nuclear genome. We compared this VGP 1.7 assembly (bTaeGut1.4) with the zebra finch VGP v1.0 pipeline (bTaeGut1.0; GCF_003957565.1) by alignment using Cactus^36^. Based on the region of false duplication we found in bTaeGut1.0, the homologous regions of false duplication were extracted by Hal^45^. We calculated the uncorrected amount of false duplications in bTaeGut1.4 from each false duplication in bTaeGut1.0 as follows: Given a length of homologous sequence *H* of a false duplication (*FD*) from new (*v1*.*7*) and prior (*v1*.*0*) VGP zebra finch in an alignment block, an uncorrected FD was calculated as *uncorrected FD* = Σ*H*_*v1*.*7*_ - (Σ*H*_*v1*.*0*_ - *FD*). If the uncorrected false duplications were ≤ 0 bp, we regarded that false duplication was corrected in the bTaeGut1.4.

## Extended Data Figures

**Extended Data Fig. 1.**
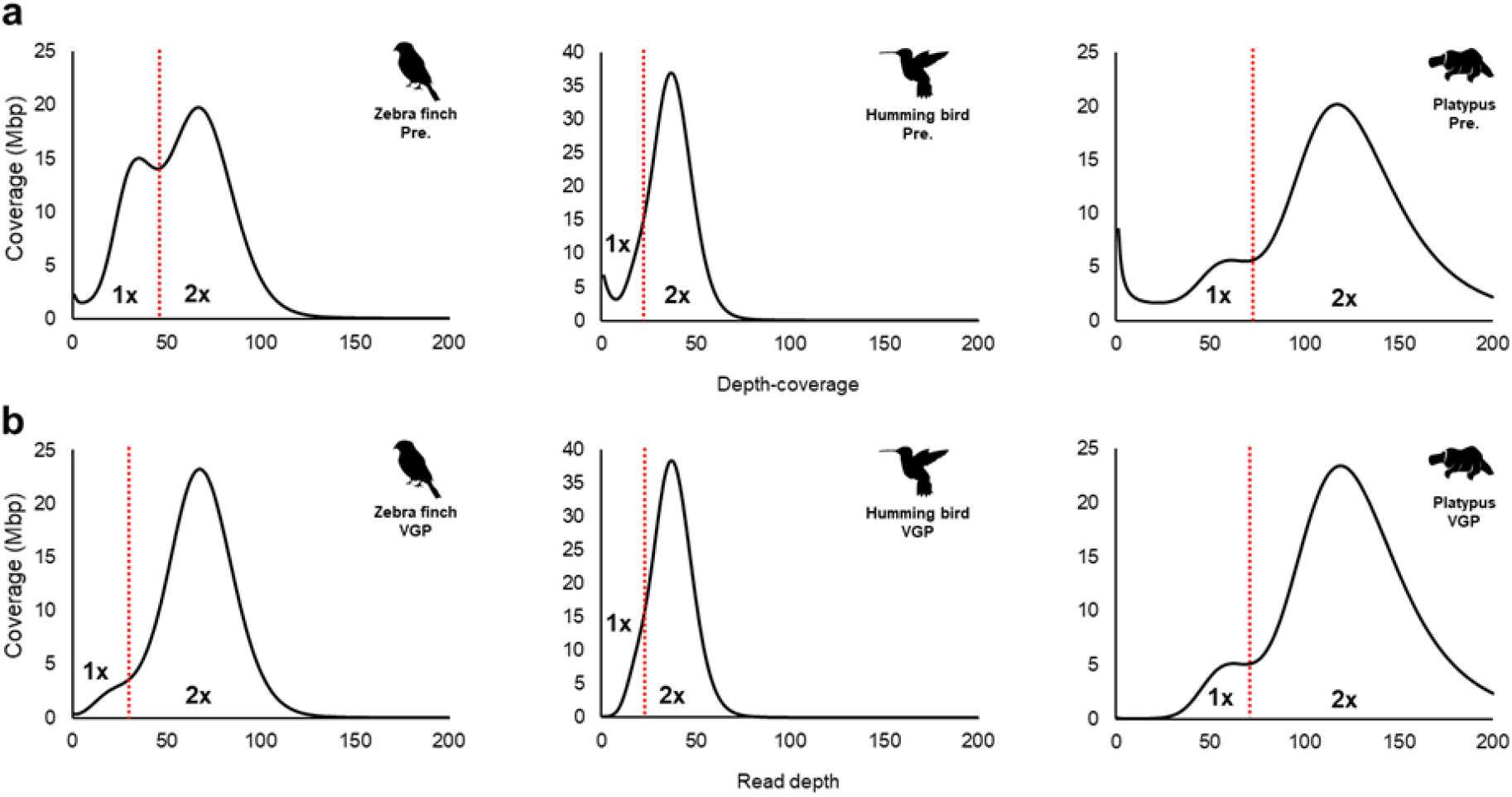
Depth-coverage profiling of all assemblies. **a**, Prior assemblies. **b**, VGP assemblies. The 10X linked-read depth-coverages of every site is summarized as a distribution. The red line shows the threshold of depth-coverage that we used to determine false duplications (to the left of the red line). Bimodal distribution in the zebra finch and the platypus assembly is caused by highly heterozygous regions.

**Extended Data Fig. 2.**
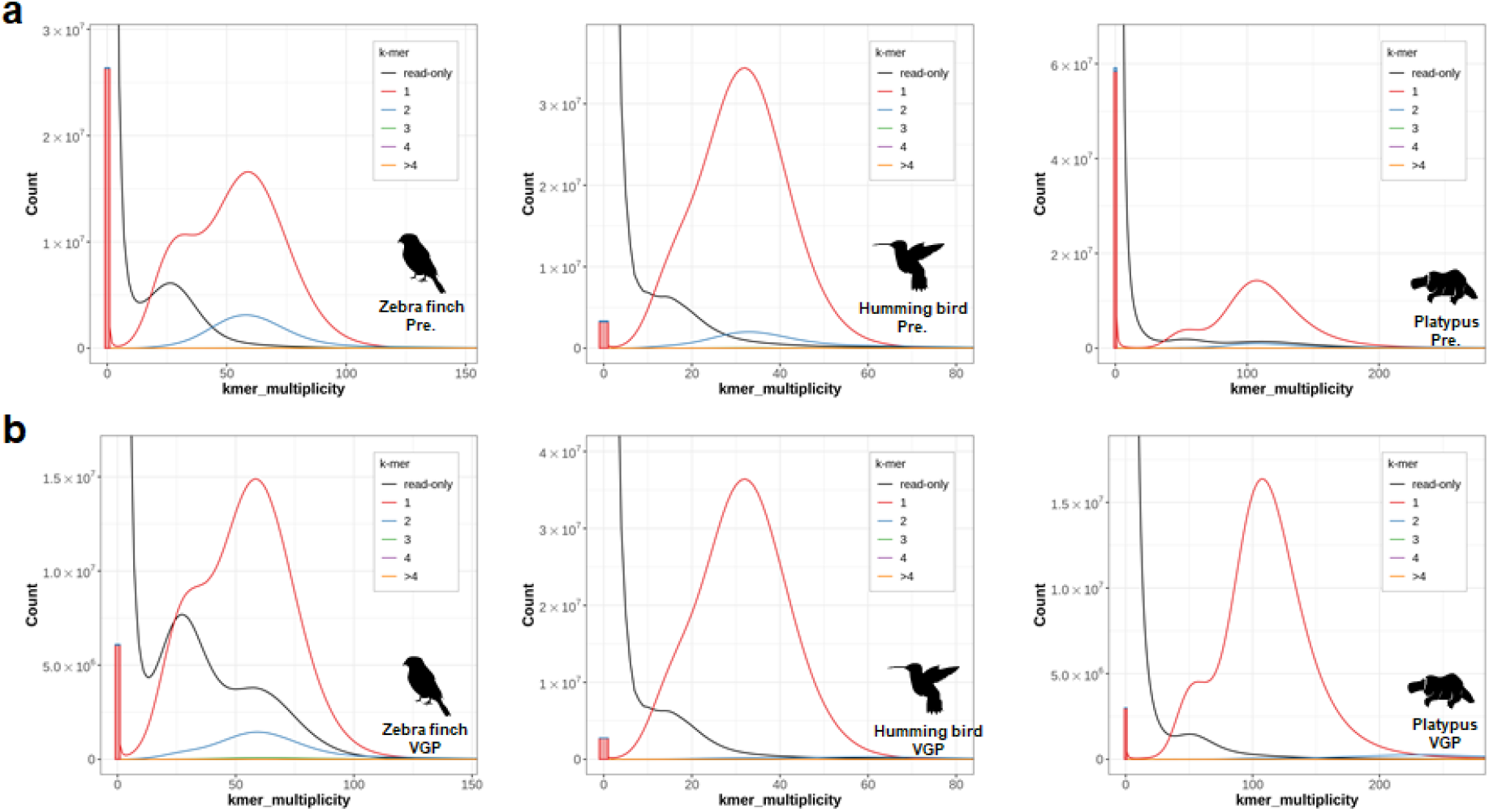
*K-mer* profiling for all assemblies. **a**, Prior assemblies. **b**, VGP assemblies. From the sequences of 10X Linked-reads and assemblies, *k-mer* multiplicity was calculated. The x-axis is the *k-mer* multiplicity calculated from reads, and the numbers in the box represents the *k-mer* multiplicity found in the primary pseudo-haplotype assembly. *K-mer* multiplicity of 2 copies or higher under the area of single copies (red) are overly represented as false duplications.

**Extended Data Fig. 3.**
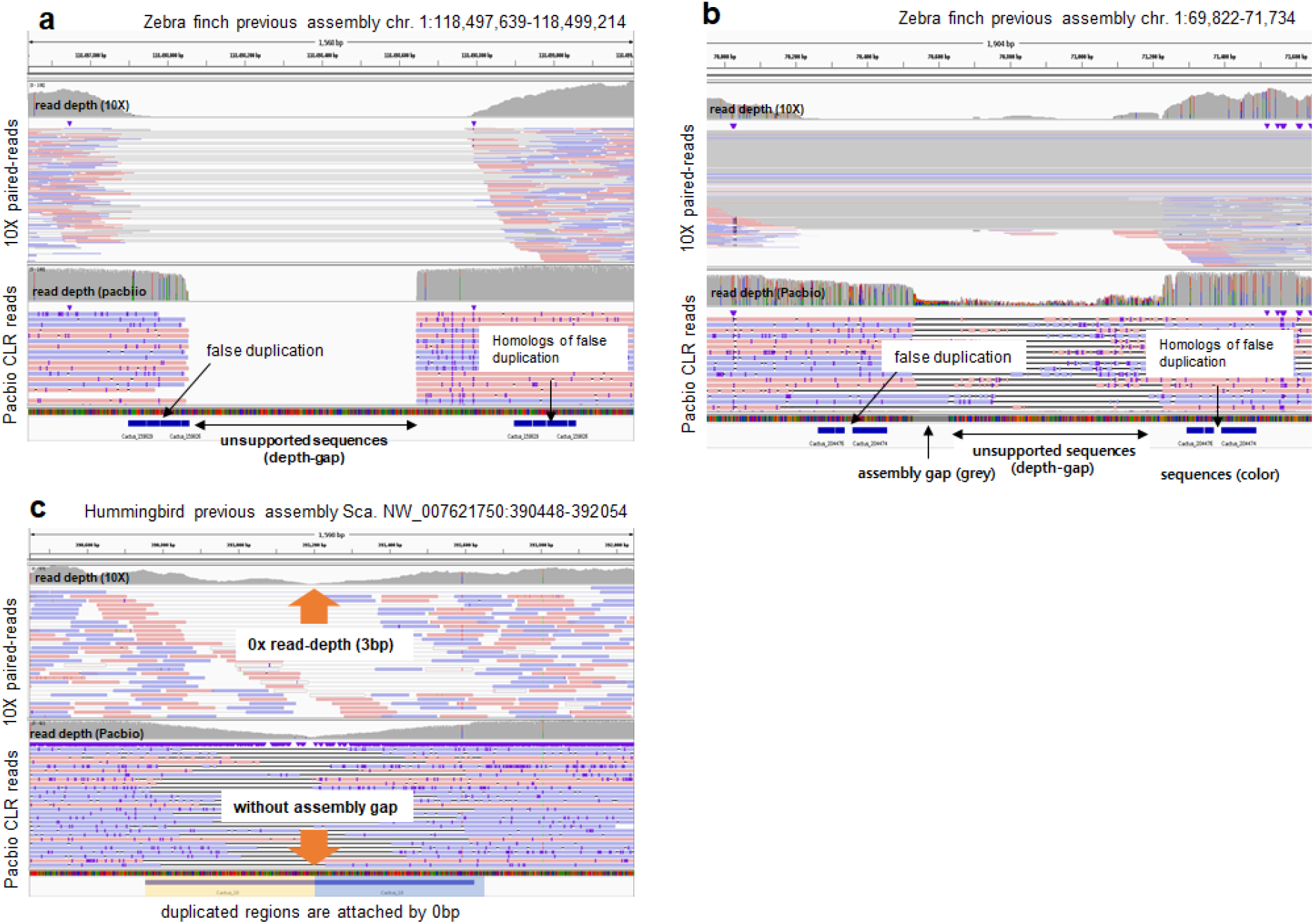
Unsupported sequences with or without assembly gaps. **a**, Unsupported sequence with a depth-gap but no assembly gap, between a false duplication. **b**, Unsupported sequence observed with an assembly gap, between a false duplication. **c**, Unsupported sequences with 0 bp in the middle between a false duplication. Unsupported sequences in the assembly were identified with 10X linked-read and Pacbio CLR alignments with no depth of coverage.

**Extended Data Fig. 4.**
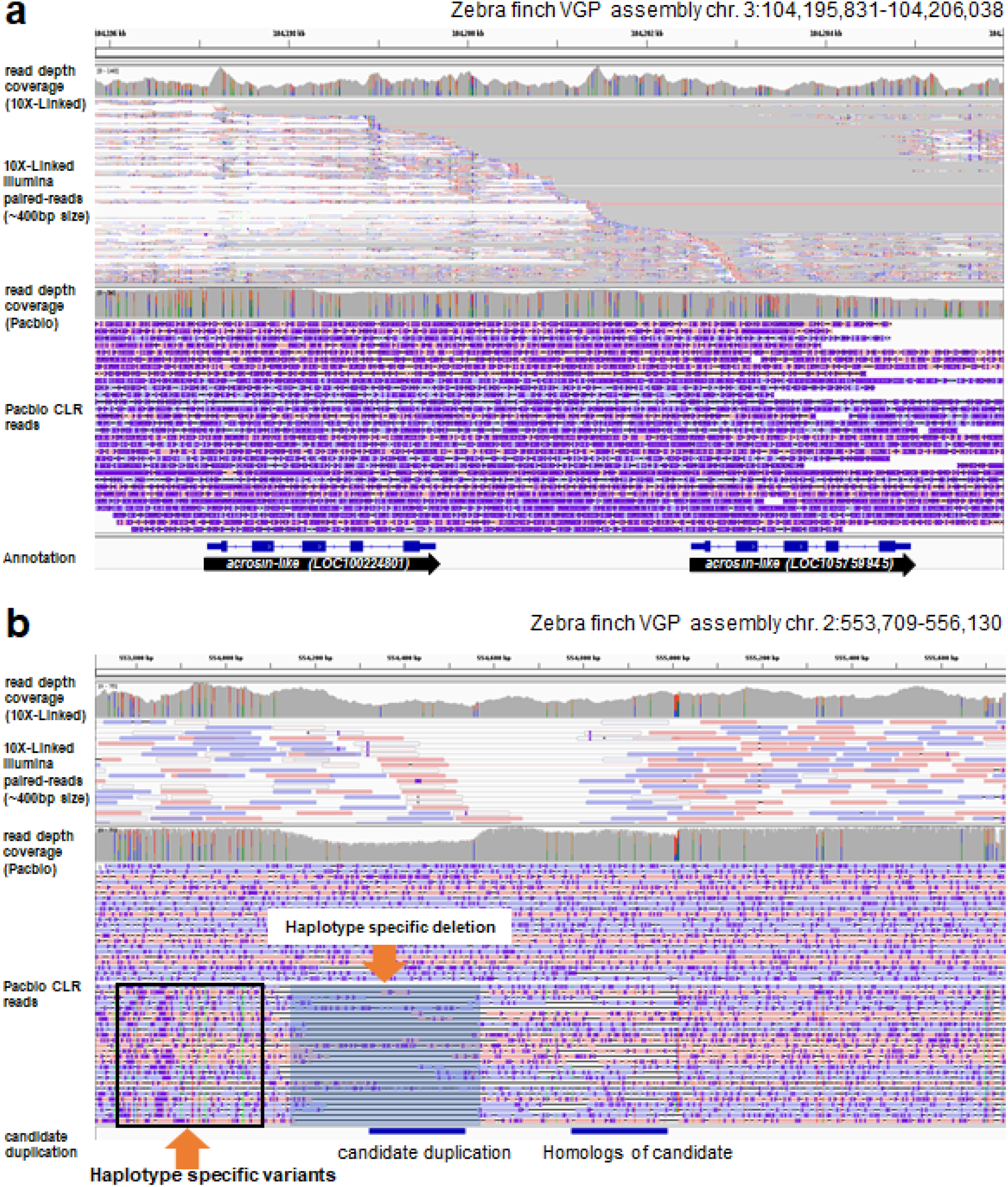
True duplication in a VGP assembly. **a**, True gene duplication of the acrosin (*ACR*) gene in the zebra finch. The 10X linked-read alignment shows discordant read alignments but has no signature of a depth-gap and decreased read-depth coverage. PacBio CLR reads alignments connect these duplicated genes with no gaps, single molecules reads, and no unsupported sequence. **b**, True haplotype specific sequence duplication with lower read depth in the zebra finch. A candidate duplication was identified as a true genomic duplication, but in one haplotype. The 10X linked-read alignment shows discordant read alignments but has no signature of a depth-gap. Half of the PacBio CLR read alignments show the allele specific duplication, while the other half show a deletion on one of the two alleles.

**Extended Data Fig. 5.**
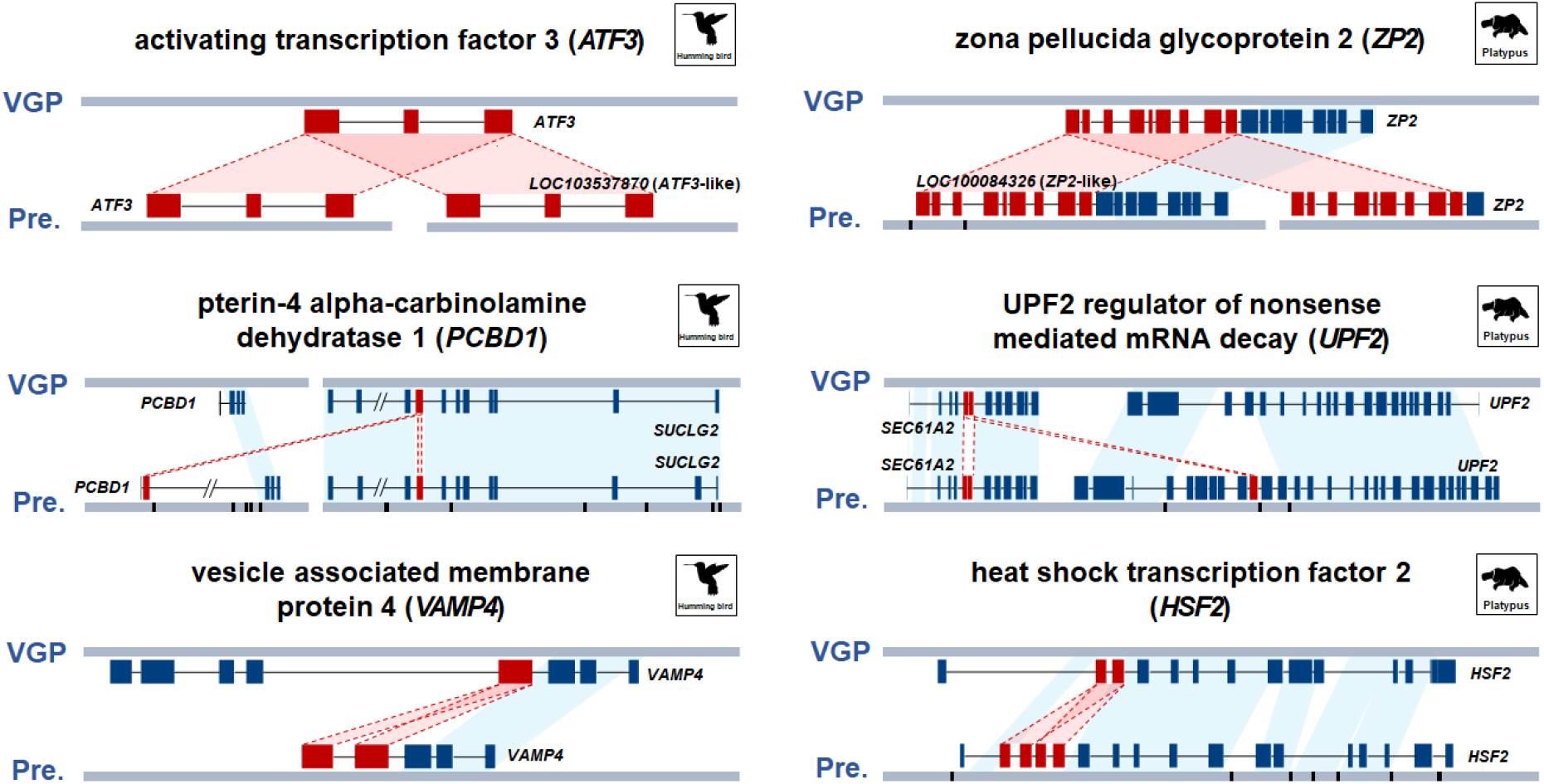
Cases of false gene gain annotations in the prior hummingbird and platypus assemblies. Top row of each alignment shows the VGP 1.0 assembly structure and annotation. Bottom row shows the previous assembly structure and annotation. The red lines represent boundaries of the false duplicated exons in the prior assemblies that are correctly assembled in the VGP assembly. The blue boxes represent the correctly assembled exons in both the previous and VGP assemblies. The black bars represent the assembly gaps on the scaffolds.

**Extended Data Fig. 6.**
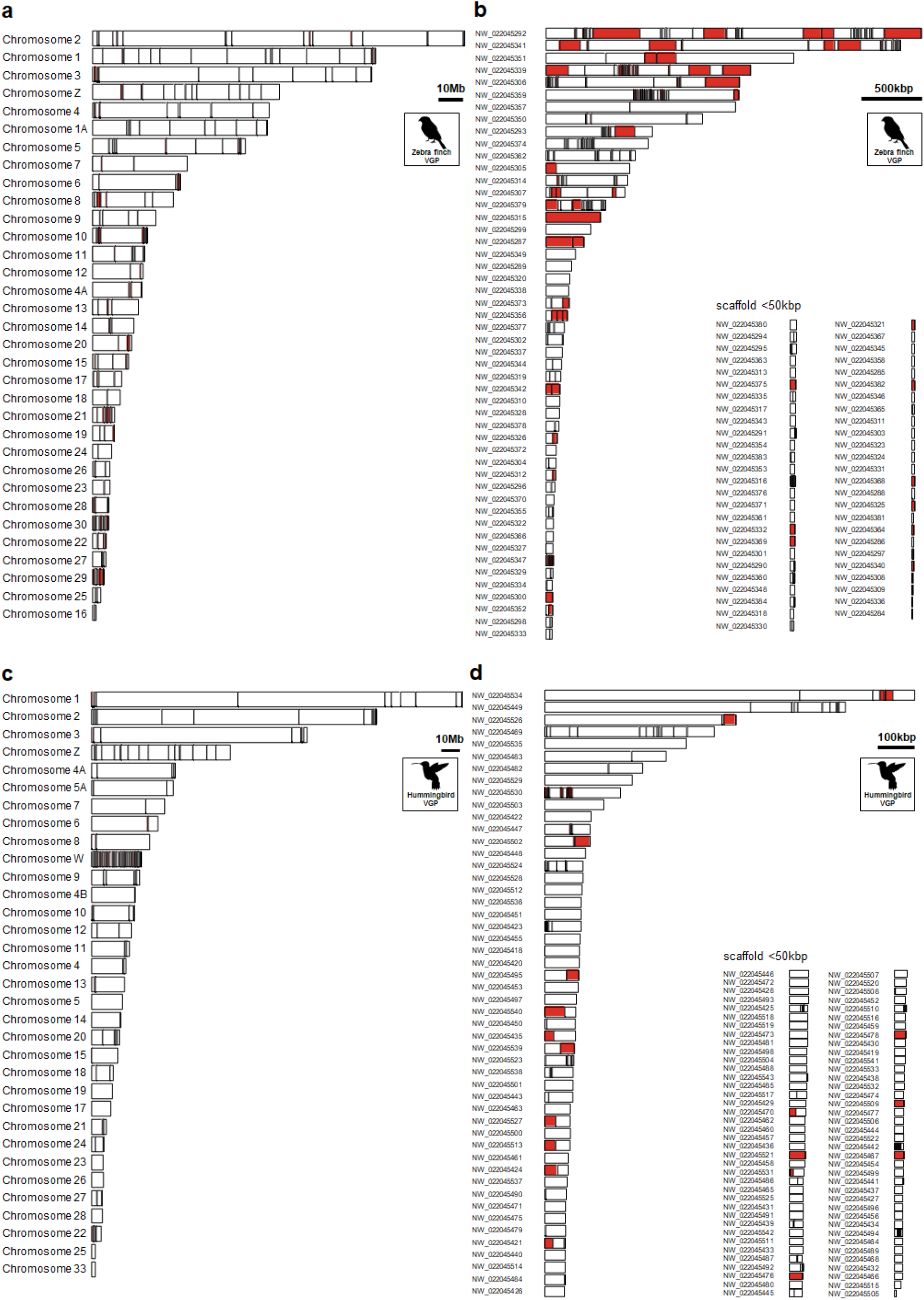
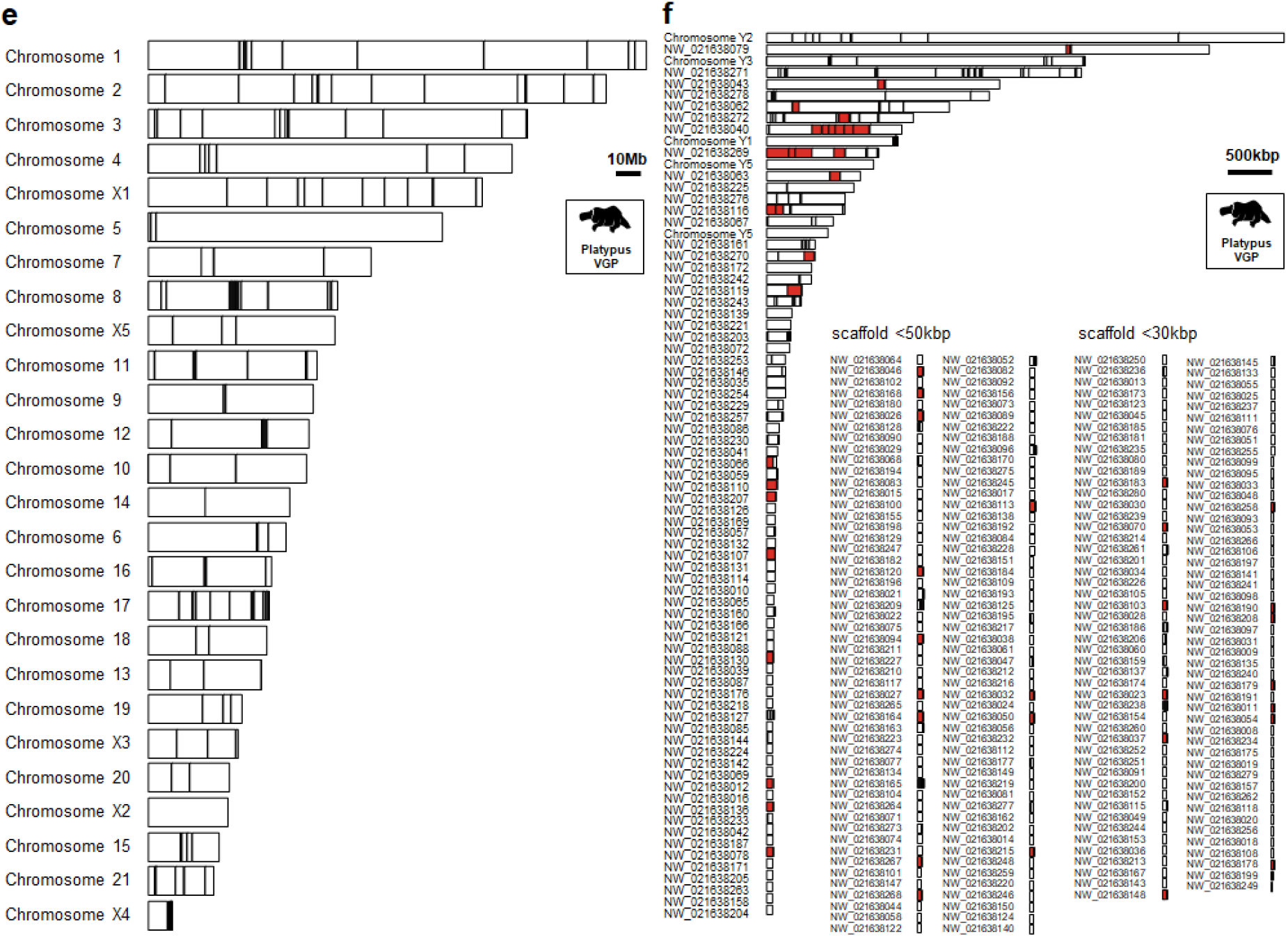
The location of false duplications in VGP assemblies. False duplications are marked as small (black, <1kbp) or large (≥ 1kbp, red) bars, in each named chromosome (**a, c, e**) or unplaced scaffold (**b, c, d**) for the zebra finch (**a, b**), hummingbird (**c, d**) and platypus (**e, f**).

**Extended Data Fig. 7.**
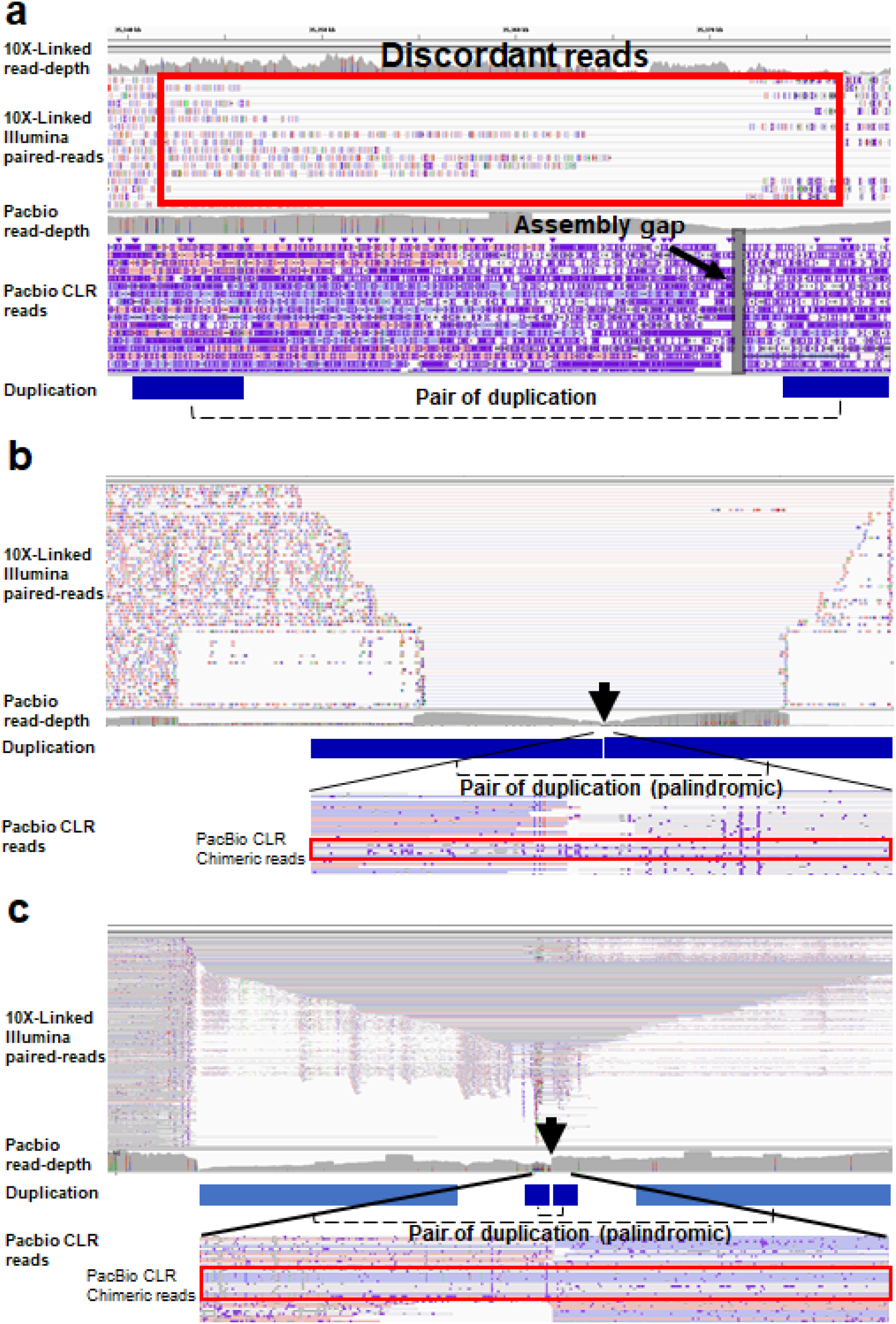
Example cases of false duplications in the VGP assemblies. **a**, A false duplicated region on zebra finch chromosome 6, ∼7 kbp long, with an assembly gap and discordant 10X linked-reads around the duplication (red box). **b**, False duplication in zebra finch scaffold NW_022045321 caused by a Pacbio sequence read chimera. The ∼10 kbp of palindromic sequence was duplicated without an assembly gap. But this region includes a sequence depth-gap with 10X linked-reads (black arrow), signifying sequencing artifacts connecting the two regions. The symmetric reduction of insert sizes of 10X linked-reads signifies the duplication of palindromic sequence. Near the center of the duplication, the six pacbio reads containing the chimeric sequences overlapped 10X depth-gap connecting the two duplicated sequences (red box). **c**, A duplicated region on chromosome 17 of the platypus, similar to the chimeric type in (a) except the 10X linked-read depth has a more pyramid structure in the palindromic duplicated regions.

**Extended Data Fig. 8.**
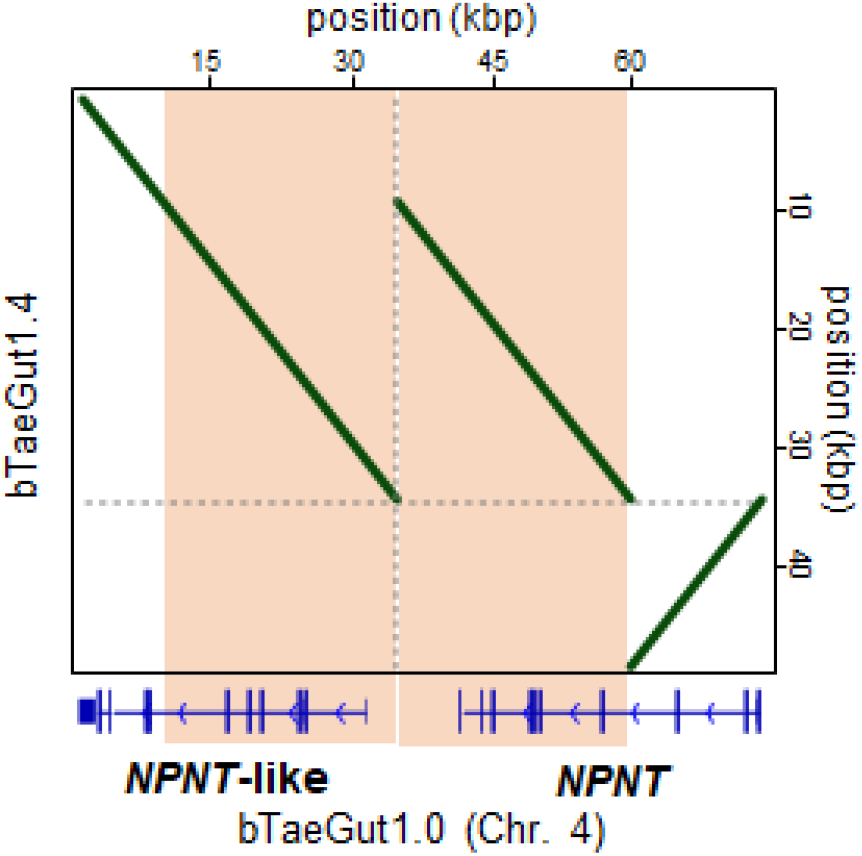
Correction of *NPNT* gene in VGP v1.7 pipeline assembly. Alignment dot-plot show that the region with six duplicated exons of *NPNT* gene in VGP v1.0 pipeline assembly (bTaeGut1.0) was prevented in VGP v1.7 pipeline assembly (bTaeGut1.4). The blue bars represent exons of *NPNT* and *NPNT-like* genes in bTaeGut1.0.

## Supplementary Information

### Supplementary Tables

**Supplementary Table 1. Statistics of previous and VGP assemblies**. Contig NG50 and Scaffold NG50 for each assembly were calculated using a source code in the VGP repository (https://github.com/VGP/vgp-assembly).

**Supplementary Table 2. Mis-annotations caused by false duplications in both previous and VGP assemblies**. The mis-annotation cases include false gene gain (FGG), false chimeric gain (FCG), and false exon gain (FEG). If the CDS overlapped caused by the false duplication was found in both -like gene and the original genes, they were named to FGG (mixed). This occurs by false duplication without haplotype phasing. Gene IDs and gene symbols are represented based on NCBI.

**Supplementary Table 3. False duplication of *V1R* family genes in the previous platypus assembly**. Only *V1R* false duplications are listed. There were a total of 43 of 44 *V1R* genes with 100% of the gene sequence was a false duplication.

**Supplementary Table 4. False duplications on transposable elements in previous assemblies**. Long terminal repeats (LTRs), short interspersed nuclear elements (SINEs), and long interspersed nuclear elements (LINEs) in each previous assembly of zebra finch, Anna’s hummingbird and platypus assemblies were searched for false duplication overlap. NCBI repeat information was used.

**Supplementary Table 5. Reduction of false duplications in the reassembled bTaeGut1.4 zebra finch genome with the VGP v1.7 pipeline**. Amount of uncorrected false duplication in bTaeGut1.4 was calculated by Σ*Hv1*.*7* - (Σ*Hv1*.*0* - *FD*), where the *H* is the length of homologous sequence of the false duplication in each assembly (v1.7 and v1.0 pipelines). A negative value in column G represents the lack of homologous sequences than expected, after false duplication correction (Σ*Hv1*.*0* - *FD*).

## Notes

### Competing Interest Statement

The authors have declared no competing interest.

